# Genetic compatibility exceeds possible ‘good genes’ effects of sexual selection in lake char

**DOI:** 10.1101/2022.03.03.482834

**Authors:** Laura Garaud, David Nusbaumer, Christian de Guttry, Lucas Marques da Cunha, Laurie Ançay, Audrey Atherton, Emilien Lasne, Claus Wedekind

## Abstract

Mating is rarely random in nature, but the effects of mate choice on offspring performance are still poorly understood. We sampled in total 47 wild lake char (*Salvelinus umbla*) during two breeding seasons and used their gametes to investigate the genetic consequences of different mating scenarios. In a first study, 1,464 embryos that resulted from sperm competition trials were raised singly in either a stress- or non-stress environment. Offspring growth turned out to be strongly reduced with increased genetic relatedness between the parents while male coloration (that reveal aspects of male health) was no significant predictor of offspring performance. In a second experiment one year later, block-wise full-factorial *in vitro* breeding was used to produce 3,094 embryos that were raised singly after sublethal exposures to a pathogen or water only. Offspring growth was again strongly reduced with increased genetic relatedness between the parents while male coloration was no significant predictor of offspring performance. We conclude that the genetic benefits of mate choice would be strongest if females avoided genetic similarity, while male breeding colors seem more relevant in intra-sexual selection.

**Impact Summary:** Males and females usually compete for access to mating partners, and they usually choose their mates. Sexual selection is therefore a major force in evolution. It shapes sexual signals and mate preferences depending on the type of mating system. A comparatively simple mating system is when fertilization is external and neither males nor females care for their brood, as is the case in salmonid fish. A group of hypotheses then predicts that female mate preferences have evolved to maximize offspring growth and survival through indirect genetic benefits. There are two types of such indirect benefits. In models of ‘good genes’ sexual selection, conspicuous signals reveal a male’s health and vigor because only males in good health can afford these costly traits. Females would then prefer males with strong signals. In ‘compatible genes’ models, females would instead focus on signals that allow them to complement their own genotype to achieve high offspring viability. An example of the latter is inbreeding avoidance through odors that reveal kinship. We sampled wild lake char to compare the likely consequences of these two types of possible female preferences for offspring growth and survival. We experimentally crossed these fish *in vitro* and raised large numbers of offspring singly and for several months. Our first experiment revealed that offspring growth would be significantly increased if females would avoid mating with genetically more similar males, while preferring males with strong sexual ornaments (in this case: yellow skin colors) would not improve offspring performance. These results could be confirmed in a second experiment with a larger sample size. We conclude that the genetic benefit of mate choice is largest if females aim for compatible genes rather than focusing on the breeding colors that males display. These breeding colors are therefore likely to play a more important role in other contexts, e.g., in male-male competition.

## Introduction

Female choice characterizes sexual selection in many species and can, in principle, happen before release of gametes (mate choice), after release of gametes and during fertilization (cryptic female choice), or after fertilization (differential investment) (Andersson 1994). When selecting mates, females can get direct benefits (nuptial gifts, protection, paternal care, etc.) or indirect (genetic) ones that increase offspring fitness. The genetic consequence of mate choice is therefore best studied in species with external fertilization and little or no parental care, and ideally where female differential investment cannot be a potentially confounding factor (Neff and Pitcher 2005). Salmonid fish are excellent models in this context because they spawn externally and leave their embryos to develop in the interstices of gravel. Moreover, much is known about their ecology and life history. Experimental fertilizations and the rearing of large numbers of embryos, each in its own container, allow separating paternal from maternal effects on offspring phenotypes and testing for various types of interactions with the necessary numbers of independent replicates.

Salmonid fish have polygamous mating systems that have been discussed as potentially influenced by intra-sexual dominance, endurance rivalry, scramble competition, sperm competition, and mate choice (Esteve 2005; Auld et al. 2019). Intra-sexual dominance is usually determined in fights or displays of fighting ability. Normally, larger males with well-developed secondary sexual characters win such dominance contests in salmonids (e.g. Jacob et al. 2007; Neff et al. 2008) and other animals (Andersson 1994). Male reproductive success is also dependent on how long they can remain reproductively active during the breeding season (endurance rivalry), how close they can position themselves to the vent of spawning females on their own and on other males’ spawning territories (scramble competition), and on their sperm number, motility, velocity, and longevity during simultaneous spawning with competitors (sperm competition). In comparison to these first four characteristics of spawning, female choice is often assumed to play a minor role in salmonid fish. However, females have frequently been observed to delay spawning when courted by non-desired males (Berejikian et al. 2000; Petersson and Jarvi 2001) or show aggressive behavior towards some types of males (Garner et al. 2010), i.e. female choice seems still possible (Esteve 2005; Auld et al. 2019).

Potential genetic benefits of mate choice can be grouped into two categories: additive genetic effects (‘good genes’) and non-additive genetic effects (‘compatible genes’) (Neff and Pitcher 2005; Achorn and Rosenthal 2020). Additive genetic effects are typically assumed to be revealed by a male’s health and vigor, because only males in good health and vigor can afford the costs that are linked to extra-ordinary colors, morphological structures, or behaviors that would potentially make males sexually attractive (Andersson 1994). The correlation between such signals of attractiveness and the signaler’s health and vigor could theoretically be strong (Fry 2022) but is expected to vary, for example, because of age-specific signaling strategies in iteroparous species (Proulx et al. 2002). In fish, additive genetic benefits have indeed been found to be strong (e.g. Wedekind et al. 2001; Pitcher and Neff 2007; Kekäläinen et al. 2010) or weak (e.g. Wedekind et al. 2008; Houde et al. 2016) and are not sufficiently understood yet.

When mate choice is driven by non-additive genetic benefits, sexual signals need not be based on costly and conspicuous traits but can instead be non-handicapping signals such as, for example, body or urinary odors (Wedekind 1994). Most examples of mate preference for genetic compatibility are about assortative mating in hybrid zones and inbreeding avoidance (Tregenza and Wedell 2000). Both types of non-additive genetic effects may be achieved in various ways, including mate choice based on genetic characteristics that would typically reveal kinship, for example genes of the major histocompatibility complex (MHC) (Ruff et al. 2012; Davies 2013; Kamiya et al. 2014).

Here we compare the potential benefits of mating with colorful males in good health and vigor (i.e., ‘good genes’ sexual selection) to the potential benefits of mating with males with complementary genotypes. We use the lake char (*Salvelinus umbla*) as model. This is a non-migratory salmonid that is endemic to lakes in the Alpine region and closely related to the Arctic char (*S. alpinus*) (Kottelat and Freyhof 2007). While Arctic char often develop strong red breeding colorations in both sexes (Janhunen et al. 2011), male lake char of our study population mostly develop a yellow breeding coloration while females remain pale (Supplementary Fig. S1). In wild Arctic char, coloration tends to increase with body length (Figenschou et al. 2013; Johansen et al. 2019), and redder males have been observed to show higher plasma testosterone levels (Johansen et al. 2019), suffer less from infections (Johansen et al. 2019), and show lower lymphocyte counts (Skarstein and Folstad 1996). Wild lake char show similar patterns: more brightly colored males are typically larger and suffer less from relative lymphocytosis and thrombocytosis, two indicators of acute infections and hence of current health and vigor, than pale males (Nusbaumer et al. 2021b). Breeding colors of wild-caught char therefore fulfill key expectations for ‘good-genes’ indicators in sexual selection (Andersson 1994), but their potential role in mate choice and/or male-male competition is not sufficiently clear yet (Nusbaumer et al. 2021b).

We sampled males and females from the wild and used their gametes in two separate breeding experiments to compare the genetic consequences of mating with a brightly colored male versus mating with genetically dissimilar males that would result in offspring of low inbreeding coefficients. We raised the resulting offspring with or without microbial stress to test whether and how male breeding colors or the genetic similarity between males and females affect offspring growth and stress tolerance.

## Methods

### Sampling and determining phenotypes and genotypes

Wild lake char were caught in December 2017 (10 males and 4 females) and December 2018 (25 males and 8 females) from Lake Geneva in gill nets, anaesthetized in a solution of 0.03% eugenol and ethanol at 1:9 ratio, photographed under standardized conditions (Fig. S1), and stripped for the gametes. These gametes were then used for 2 different types of fertilization experiments as explained below.

Male breeding coloration (that is mostly yellow in the study population) was analyzed from the photos in Fiji as described in (Nusbaumer et al. 2021b). Briefly, the white balance was standardized based on the white and black sections of the color scale on each photo (Fig. S1). Male yellowness was then measured as the *b\** components of the CIE-*L*a*b\** color space model (León et al. 2006), where the L* axis quantifies lightness from dark to light, the a* axis ranges from green to red, and the b* axis from blue to yellow, with high b* values indicating strong yellow color (the axes scale from -100 to 100). After transformation of the image in a Lab-Stack with the function run(“Lab Stack”), all CIE-*L*a*b\** color components were measured from 3 squares in the pectoral region, the ventral region, and the anal region (Fig. S1) and then averaged. Lightness and redness were significantly correlated with yellowness (the yellower the lighter and/or the redder (Nusbaumer et al. 2021b)) and were therefore ignored for further analyses.

The DNAeasy Blood & Tissue kit (Qiagen, Hilden, Germany) was used to extract DNA from up to 20 mg of fin samples (with an extraction robot, following manufacturer’s instructions). DNA concentration was measured using Qubit 2.0 (Thermo Fisher Scientific, Waltham, Massachusetts) while its integrity was verified on a 1% agarose gel. The DNA of each individual was standardized to a concentration of 20 ng/µl. The libraries were prepared as follow: the enzymes EcoRI-HF and MspI (New England Biolab, Ipswich MA, USA) were used for DNA digestion and a unique EcoRI barcode was ligated to each individual (Brelsford et al. 2016). After library purification, PCR amplification was performed and fragments size-selected in between 400-550bp. The parents of the first experiment were then single end genotyped on 2 lanes and the parents of the second experiment on one lane of Illumina HiSeq 2500 with fragments of 150 bp length at Lausanne Genomic Technologies Facility (University of Lausanne).

The resulting ddRAD sequencing data of the two experiments were analyzed separately using Stacks (Catchen et al. 2013) with the default parameters unless otherwise specified. Individuals sequences were demultiplexed using *process_radtags* (Catchen et al. 2013) in the 1^st^ experiment (see also Nusbaumer et al. 2021b) and trimmomatic (Bolger et al. 2014) in the 2^nd^ experiment and trimmed to 110bp before mapping to the *Salvelinus spp.* reference genome (*S. alpinus* and/or *S. malma*, Christensen et al. 2018) using BWA (Li and Durbin 2010). *Populations* (Catchen et al. 2013) was used to generate the VCF file considering only loci present in at least 80% of the individuals and marker heterozygosity of 0.5. To reduce incorrect heterozygosity call and remove paralogs, vcftools (Danecek et al. 2011) was used filtering the loci for a coverage between 10X and 50X for the 1^st^ experiment and between 7X and 60X in the 2^nd^ experiment, presence in all individuals, and at Hardy-Weinberg equilibrium. This led to a total of 4,150 SNPs with a mean coverage of 29X for all individuals of the 1^st^ experiment, and a total of 975 SNPs with a mean coverage of 27X in 30 of the 33 individuals of the 2018 samples. Three males with low genotyping rate (>30% missing data) were excluded from all further analyses. Individual inbreeding coefficients (F_β.parent_) and kinship (µF_β.offspring_) were estimated for each breeding experiment separately, using the *beta.dosage* function of the package *Hierfstat* 0.04-30 (Goudet 2005).

### Experiment 1

The first experiment was done with the 14 lake char that were sampled in 2017. Milt was stripped from the males into individual 50 mL Falcon tubes (Greiner Bio-One, Austria), carefully avoiding contamination by water, urine, or blood, then mixed 1:9 with diluted Storefish© (IMV Technologies, France; a media for preservation and dilution of fish milt, here diluted with 1:9 with MilliQ water) and stored on ice. Females were kept in a 1,000 L circular tank with free-flowing lake water for one day before eggs were stripped into individual plastic containers. The ovarian fluid each was separated from the eggs with a syringe and stored in 50 mL tubes at 4°C. The eggs were twice with Ovafish (IMV Technologies, France; a solution for washing eggs after stripping) up to 200 mL. They were then stored in approximately 20 mL of the same medium less than one hour at 4 °C to avoid drying. Sperm concentration and motility was determined for each milt sample to allow preparing standard dilutions (details in Nusbaumer et al. 2021b). Diluted milt of 2 males each (haphazardly assigned from the 10 males) was mixed such that each male was represented with 25 million active sperm per mL of the mix. One mL of such a mix was then used to fertilize approximately 24 eggs of a female in wells of 6-well plates (Falcon, BD Biosciences, Allschwil, Switzerland). This 1 mL of mix was activated in a separated tube with either 4 mL of chemically standardized water (i.e. reconstituted according to the OECD guidelines (OECD 1992)) or 4 mL of standardized water with ovarian fluid (ratio ovarian fluid to water = 1:2) and vortexed for 5s before being poured in a well with the eggs. Two minutes later, 16.8 mL standardized water was added to fill the wells. The eggs were then left undisturbed for 2 hours of egg hardening. Each mix of sperm was used full-factorial, fully balanced, and repeatedly with or without ovarian fluids on eggs of all females, resulting in a total of 80 batches of about 24 eggs each (5 types of sperm mixes x 4 females x 2 experimental conditions x 2 replicates = 80 batches). The 2 experimental conditions were created to test for potential effects of ovarian fluids on sperm competition as reported elsewhere (Nusbaumer et al. 2021b). The focus here is on offspring performance relative to fathers’ yellowness and the kinship coefficient between the parents.

Ten of these 80 batches of eggs were accidentally lost (all from the same female and exposed to sperm activated in water only), reducing the number of eggs to be monitored to N = 1,642. These eggs were batch-wise rinsed in a sterilized tea strainer under cold tap water for 30 s (4 L/min), then distributed singly to 24-well plates (each embryo in its own compartment filled with 1.8 mL autoclaved standardized water) and raised at 4.5°C (2009). The rinsing of the eggs was considered necessary to avoid very high embryos mortalities (C. Wedekind, unpublished observations), but it was done such that remainders of ovarian fluids could still be expected on the developing eggs. Apart from this organic material that was expected to support microbial growth (2010), no other environmental stress treatment was applied. The rate of embryonated eggs was determined at 28 dpf (days past fertilization). Towards the end of the embryo stage, all wells were checked daily to record embryo mortality and hatching date. Freshly hatched larvae were immediately transferred to new 5mL wells with standardized water only. Standard photos were taken on the same day to measure larval length in ImageJ (Fig. S2a). Larval length and yolk sac volumes were measured at the day of hatching and 14 days later (Fig. S2b) when larvae were euthanized and genotyped for 3 to 6 microsatellite markers to identify their fathers (details in Nusbaumer et al. 2021a). All embryos and larvae that had died before were also genotyped. A sex-specific marker was added to the first multiplex of 3 microsatellite markers to test for the sex-specific effects on embryo development that are reported in Nusbaumer et al. (2021a).

### Experiment 2

The gametes of the 25 males and 8 females of the 2018 sample were stripped into individual containers as in the first experiment. The eggs of each female were about equally distributed to 25 Petri dishes (Greiner Bio-One, Austria). Undiluted milt of one male each was added to the individual batches of eggs to produce all possible 200 families (8 females x 25 males). Standardized water was then added to activate the sperm and induce fertilizations. After letting the freshly fertilized eggs harden for 2 hours, 72 eggs per sib group (14,400 in total) were transported on ice to a climate chamber where they were batch-wise rinsed in cold tap water (in sterilized tea strainers for 30 seconds at 2 L/minute) and singly distributed to 24-well plates for two separate experiments: 24 eggs/sib group were used for the present study and the remaining 48 eggs/sib group for a study on the evolutionary potential of the study population to adapt to a common micropollutant (Garaud et al. in preparation). The rate of embryonated eggs was determined at 35 dpf.

At 69 dpf, half of the eggs per sib group were exposed to the bacterium *Aeromonas salmonicida* while the other half was sham treated. This treatment was prepared as follows: Dry-frozen *A. salmonicida* that had previously been isolated from brown trout (*Salmo trutta*) were rehydrated and inoculated at 22°C for 24 hours in TBS (Tryptic Soy Broth, Fluka, Switzerland). Cultures were then washed and counted in a Helber counting chamber. Bacteria were diluted in autoclaved standardized water and 1% TSB such that adding 100 µL in each well plus 100 µL MilliQ water would result in 0.5 x 10^6^ cfu/mL in the well. The sham treated embryos received 200 µL standardized water. The bacterial density was chosen to be likely sublethal (D. Nusbaumer, unpublished observations) so that the more informative indicators of embryo development rate could be used as dependent variable instead of the binary embryo mortality.

Towards the expected start of hatching, all wells were checked daily to record embryo mortality and hatching date. Freshly hatched larvae were immediately transferred to new 5mL wells with standardized water only. Standard photos were taken on the same day to determine larval length in ImageJ.

Because larval size measurements turned out to be statistically linked to the timing of hatching (see Results), larval length was again measured 14, 28, and 42 dph (days past hatching) while yolk sac volume was determined at the day of hatching and at 42 dph. From these measurements, larval length at 130 dpf (L_130dpf_), a haphazardly chosen time point that was defined by the time of fertilization (instead of the time of hatching) and that was shortly before the last length measurement (Fig. S2, S3) was calculated as

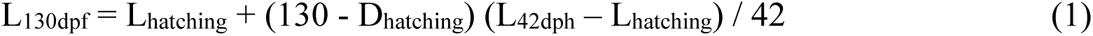

where L_hatching_ is the larval length at hatching, D_hatching_ the number of days from fertilization to hatching, and L_42dph_ the larval length at 42 dph. Yolk sac volume at 130 dpf was determined as

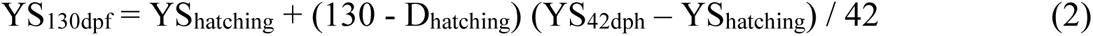

where YS_hatching_ is the yolk sac volume at hatching and YS_42dph_ the yolk sac volume at 42 dph.

### Statistics

Statistical analyses were done in JMP 15.2.1. When using linear mixed models, family identity and their interactions with treatments were included as random factors while treatment, male yellowness or genetic distance between males and females (kinship) were used as fixed factors. For size measurements as dependent variables, hatching date was included as further fixed factor.

## Results

### Experiment 1

The F_β.parent_ of the 14 males and females was on average 0.056 (± 0.025 SD) and ranged from 0.003 to 0.093. Since all F_β.parent_ were within 2 SDs from the mean, all 40 families could be included in the statistical analyses.

Table 1 gives the effects of male yellowness and kinship between males and females on measures of offspring growth while controlling for the family and treatment effects that have been reported before (Nusbaumer et al. 2021a). Male yellowness did not significantly predict any offspring characteristics (Table 1a; Table S1a; Fig. 1a-c, Fig. S4a,b). However, the genetic distance between males and females (kinship) was a strong predictor of embryo and larval growth: With increasing kinship, the larvae hatched at smaller length and with smaller yolk sacs despite hatching at similar times (Table 1b; Fig. 1d-f). Fourteen days after hatching, the effects of kinship on larval length were no more statistically significant, but yolk sac volumes were still smaller with increased kinship (Table S1b; Fig. S4c,d).

**Figure 1.**
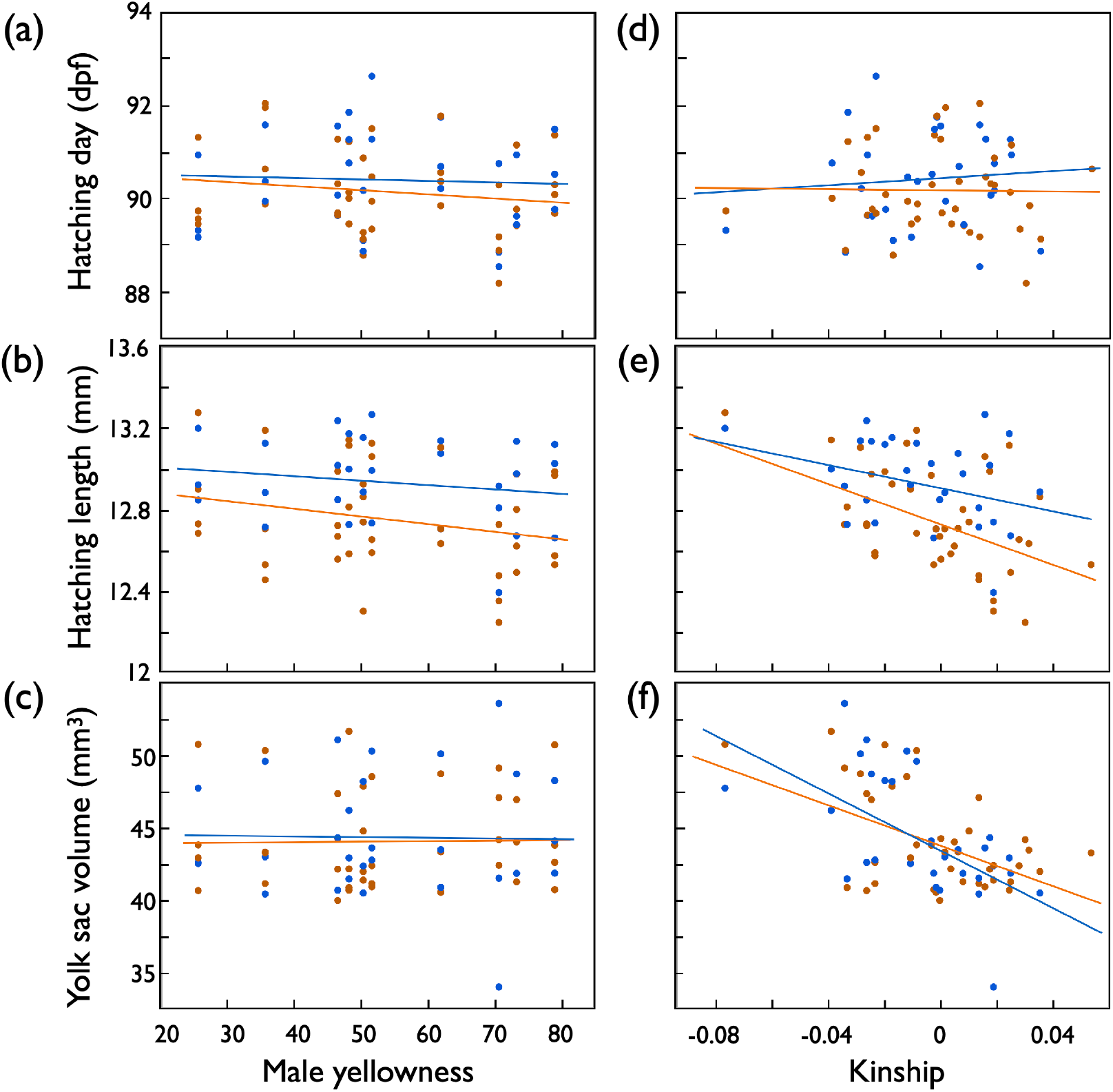
First experiment (fertilization during sperm competition): hatching time (dpf, days past fertilization), hatchling length, and yolk sac volume of hatchlings (means per full-sib family) predicted by (a,b,c) male skin coloration (yellowness), (d,e,f) the genetic similarity between males and females (kinship) after exposure to organic pollution (orange symbols and regression lines), or not exposed (blue). See Table 1 for statistics.

**Table 1.** First experiment: Linear mixed model (REML, unbounded variance components) on embryo hatching time (dpf) and hatchling length when predicted by treatment and (a) male skin coloration (yellowness) or (b) the genetic similarity between males and females (kinship). Hatching date was included as further fixed factor when testing for effects on hatchling length. Family identity and the interaction between family ID and treatment were included as random factors. Significant p-values are highlighted in bold.

| Effects | d.f. | Hatching time |  |  | Hatchling length |  |  | Yolk sac volume |  |  |
| --- | --- | --- | --- | --- | --- | --- | --- | --- | --- | --- |
|  |  | F | Variance component | p | F | Variance component | p | F | Variance component | P |
| *(a) Male skin coloration* |  |  |  |  |  |  |  |  |  |  |
| Fixed effects: |  |  |  |  |  |  |  |  |  |  |
| Treatment | 1 | 0.3 |  | 0.56 | 19.0 |  | <0.001 | 0.1 |  | 0.74 |
| Yellowness | 1 | 0.4 |  | 0.53 | 0.9 |  | 0.36 | 0.01 |  | 0.93 |
| Treatment x yellowness | 1 | 0.4 |  | 0.54 | 0.4 |  | 0.54 | 0.02 |  | 0.90 |
| Hatching date | 1 |  |  |  | 36.6 |  | <0.001 | 7.8 |  | 0.005 |
| Random effects: |  |  |  |  |  |  |  |  |  |  |
| Family ID<sup>1</sup> |  |  | 0.70±0.20 | <0.001 |  | 0.05±0.01 | <0.001 |  | 10.4±2.8 | <0.001 |
| Family x treatment<sup>1</sup> |  |  | 0.12±0.06 | 0.04 |  | 0.004±0.004 | 0.26 |  | 0.5±0.6 | 0.42 |
| Residual |  |  | 2.18±0.09 |  |  | 0.16±0.01 |  |  | 35.0±1.4 |  |
| *(b) Genetic similarity* |  |  |  |  |  |  |  |  |  |  |
| Fixed effects: |  |  |  |  |  |  |  |  |  |  |
| Treatment | 1 | 0.3 |  | 0.59 | 20.9 |  | <0.001 | 0.02 |  | 0.89 |
| Kinship | 1 | 0.003 |  | 0.96 | 7.6 |  | 0.009 | 15.0 |  | <0.001 |
| Treatment x kinship | 1 | 0.02 |  | 0.90 | 3.0 |  | 0.09 | 0.04 |  | 0.84 |
| Hatching date | 1 |  |  |  | 35.5 |  | <0.001 | 6.7 |  | 0.01 |
| Random effects: |  |  |  |  |  |  |  |  |  |  |
| Family ID<sup>1</sup> |  |  | 0.70±0.20 | <0.001 |  | 0.04±0.01 | <0.001 |  | 6.9±2.0 | <0.001 |
| Family x treatment<sup>1</sup> |  |  | 0.12±0.06 | 0.04 |  | 0.003±0.003 | 0.42 |  | 0.5±0.7 | 0.47 |
| Residual |  |  | 2.18±0.09 |  |  | 0.16±0.01 |  |  | 35.0±1.4 |  |

<sup>1</sup>REML unbounded variance components ± standard error, Wald p-values

### Experiment 2

The mean F_β.parent_ of the 30 males and females that could be genotyped was 0.006 (± 0.092 SD). One male had an F_β.parent_ of -0.328 that was > 3 SDs smaller than the overall mean. It was therefore considered an outlier and excluded from all analyses, in addition to the three males that could not be genotyped. All other F_β.parent_ were within 2 SDs from the mean, i.e., the embryos considered for the analyses stem from 21 males crossed with 8 females (168 families).

The overall rate of embryonated eggs at 35 dpf (before treatment was applied) was 81.0% and varied between females (from 42.2% to 92.7%; χ^2^ = 516.2, df = 7, p < 0.001) and males (χ^2^ = 68.3, df = 20, p < 0.001). This rate could not be predicted by male yellowness (r = 0.05, n = 21, p = 0.83) nor by the mean genetic distance between a male to all females (mean kinship per male; r = 0.02, n = 21, p = 0.93). Total embryo mortality (from 35 dpf until the day of hatching) was 0.9% and not linked to treatment (χ^2^ = 0.12, df = 1, p = 0.73).

Exposure to the pathogen reduced embryo and larval growth because it not only led to earlier hatching of smaller larvae, but these larvae were also smaller than sham exposed ones at 130 dpf, i.e., at a late larval stage that was defined by time since fertilization instead of time of hatching (Table 2; Fig. 2). There were also significant family effects on embryo development rates as measured by hatching date, hatchling length, and larval size at 130 dpf (see family effects in Table 2). Mean hatching date per family was treatment dependent while hatchling length and larval size at 130 dpf were not (family x treatment interactions in Table 2).

**Figure 2.**
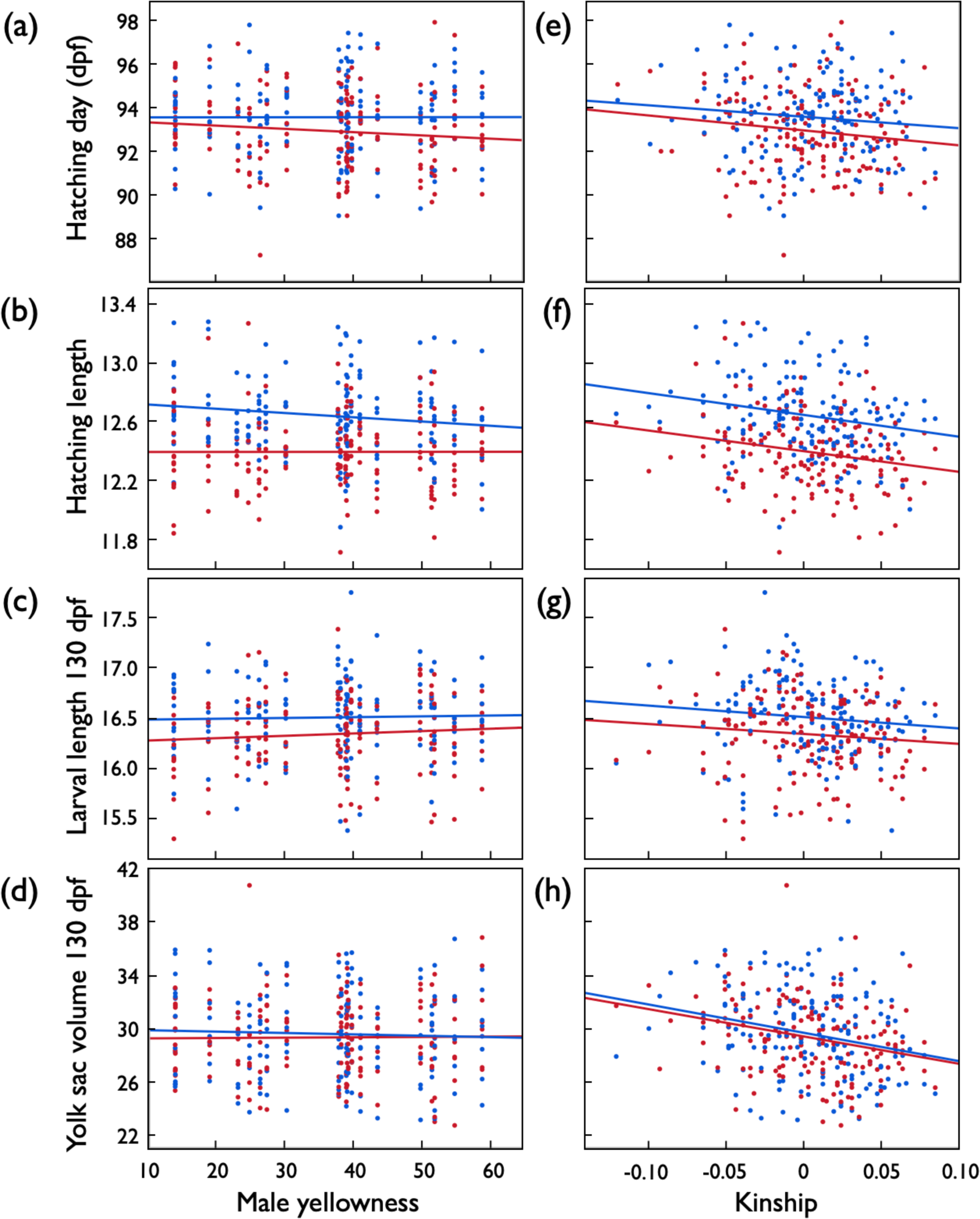
Second experiment: Embryo development after exposure to *Aeromonas salmonicida* (red symbols and regression lines) or sham exposure (blue). (a) Mean hatching time (dpf = days past fertilization), (b) mean larval length at hatching (mm), (c) mean larval length 130 dpf (mm), and (d) mean yolk sac volume 130 dpf (mm^3^) per full-sib family predicted by male skin coloration (yellowness), and (e,f,g.h) predicted by the genetic similarity between males and females (kinship). See Table 2 for statistics.

**Table 2.**
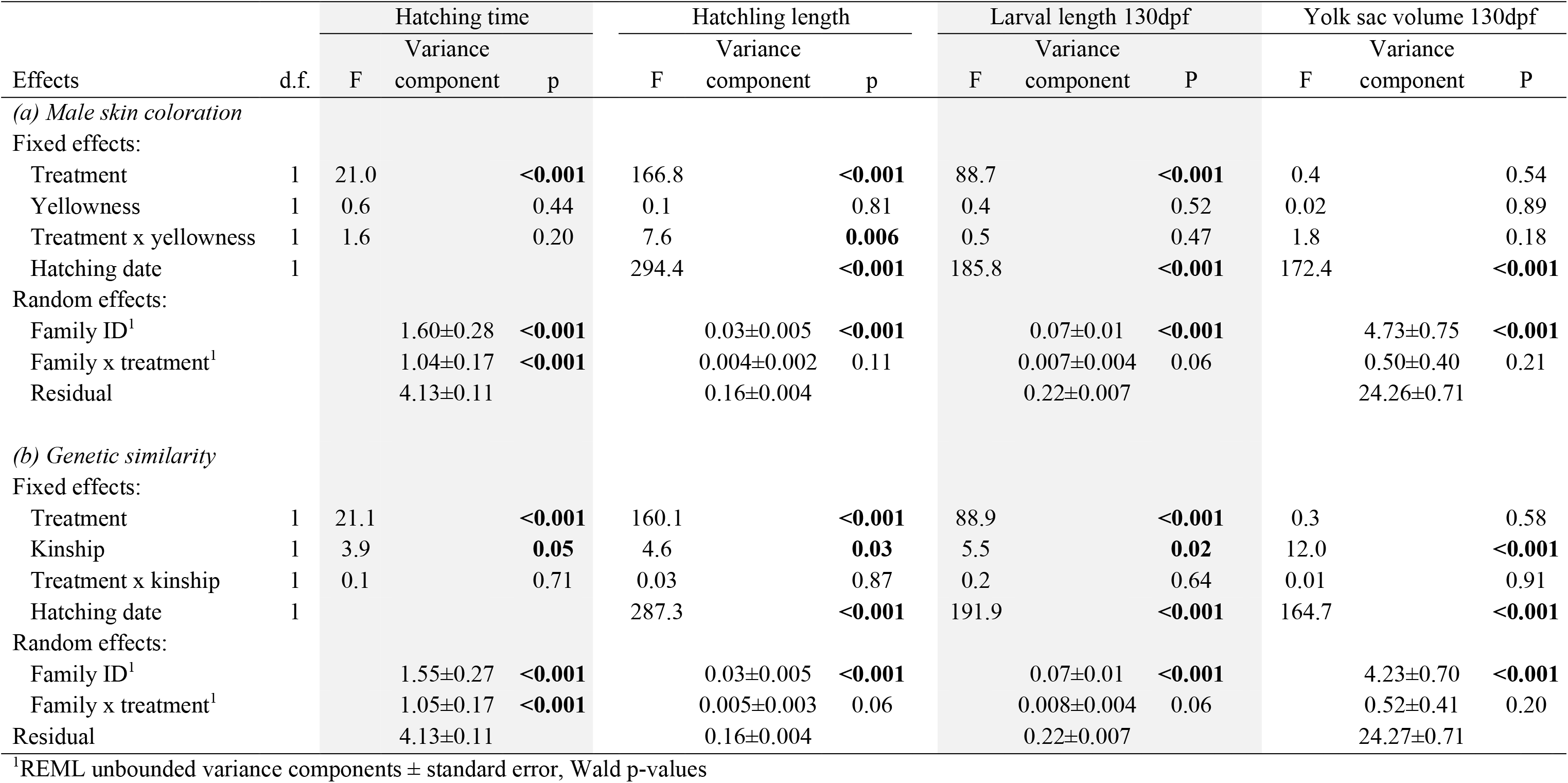
Second experiment: Linear mixed model (REML, unbounded variance components) on embryo hatching time (dpf) and hatchling length when predicted by treatment and (a) male skin coloration (yellowness) or (b) the genetic similarity between males and females (kinship). Hatching date was included as further fixed factor when testing for effects on hatchling length. Family identity and the interaction between family ID and treatment were included as random factors. Significant p-values are highlighted in bold.

Male yellowness did again not predict embryo development and growth: Neither the timing of hatching, nor hatching length or larval length at 130 dpf were correlated to male yellowness (Table 2a; Fig. 2a-d). A significant interaction term between treatment and yellowness (Table 2a) suggested that offspring of yellower males tend to hatch at smaller size when not stressed while there seemed to be no such link to male yellowness when embryos were raised under stress conditions (Fig. 2b). However, this interaction could not be confirmed at 130 dpf (Table 2a; Fig. 2c,d).

The genetic distance between males and females (kinship) was again a strong predictor of embryo and larval growth: With increasing kinship, the larvae not only hatched earlier and at smaller length and with smaller yolk sacs (Table 2b; Fig. 2e,f), but larvae at 130 dpf were increasingly smaller with decreasing genetic distance between the parents (Table 2b; Fig. 2g,h). These kinship effects on embryo and larval development were similar in both treatment groups (non-significant interaction terms in Table 2b).

### Heritability of inbreeding coefficients

The mean expected inbreeding coefficients of each male’s offspring (µF_β.offspring_) differed between years (estimates were higher in 2018 than in 2017) and could be predicted by the inbreeding coefficients of their fathers (F_β.male_): high F_β.male_ led to high µF_β.offspring_ (linear regression after removal of non-significant interaction term, intercept: t = 2.4, p = 0.02; effect of year: t = -2.6, p = 0.02; effect of F_β.male_: t = 5.2, p < 0.0001; Fig. S4).

## Discussion

We studied the growth and survival of about 4,500 singly raised embryos and larvae of 208 experimental families to test what type of female choice would provide more genetic benefits: a preference for a potential ‘good genes’ indicator (aiming at additive genetic effects) or a preference for some type of compatible genes (aiming at non-additive genetic effects). We focused on breeding coloration as potential ‘good genes’ indicator and found that this secondary sexual trait was no useful predictor of offspring performance. However, the genetic similarity between the parents (kinship) had strong effects on offspring performance: higher kinship caused significant reductions in offspring growth. These effects were consistent in two different types of experiments done in two different years.

Arguably, a study on possible genetic benefits has to be based on potential fitness indicators, but fitness is notoriously difficult to measure. Here we focused on embryo and larval growth under stress and non-stress conditions, keeping the environmental stress at sub-lethal level to avoid mortalities and to focus on the continuous and hence more informative measures of growth. Bylemans et al. (2022) recently tested the ecological relevance of such growth measurements and found that they were good predictors of juvenile size after a first spring and summer in the wild. Juvenile size has long been known to positively affect survival and reproduction (Garcia de Leaniz et al. 2007). Embryo and larval growth are therefore useful indicators of offspring fitness. We conclude that if females can choose their mate, they should avoid genetic similarity and ignore male coloration to achieve the highest genetic benefit from mate choice. Analogous negative effects of genetic similarity could recently be demonstrated in an amphibian (Byrne et al. 2021), and analogous non-significant effects of male coloration on offspring viability were recently reported for three-spined stickleback (*Gasterosteus aculeatus*) (Chiara et al. 2022).

It has been suggested that the ‘good genes’ hypothesis may be better described as ‘absence of bad genes’ because condition-dependent ornaments may reveal the absence of deleterious mutations (Tomkins et al. 2004; Baur and Berger 2020). Classical non-additive genetic effects such as inbreeding can even contribute to additive genetic variation for fitness because of genetic drift in finite populations (Neff and Pitcher 2008). Such a heritability of inbreeding coefficients is expected to be highest in small or fragmented populations (Neff and Pitcher 2008). We previously found no such statistically significant heritability when testing within the 10 males of the 2017 sample (Nusbaumer et al. 2021b). However, when combining the 2017 samples with the new sample taken in 2018 we now find that F_β.male_ is in fact significantly correlated to the average kinship coefficients per full-factorial breeding and hence to the µF_β.offspring_, i.e., we observed the heritability of inbreeding coefficients that is predicted for small populations.

We have demonstrated before that the variation in F_β.male_ in our study population was not linked to male yellowness (Nusbaumer et al. 2021b). Male yellowness was, however, positively linked to male size and measures of milt quality and of general health and vigor (Nusbaumer et al. 2021b). Male coloration in lake char may therefore first be an indicator of current fighting ability (or willingness to fight) and hence of dominance in male-male competition (Johnstone 1997). In salmonids, the capacity of a male to dominate others seems to be an important predictor of male mating success in the wild (Auld et al. 2019). Our findings suggest, however, that females get little genetic benefits from mating with such males. This supports analogous results from another salmonid (Jacob et al. 2007). It is possible that females benefit from the protection that dominant males sometimes provide against egg predation during the first minutes after spawning (Frye et al. 2021) or from higher fertilization success due to milt and sperm traits linked to male dominance (Masvaer et al. 2004; Nusbaumer et al. 2021b). However, if females aim for genetic benefits, we predict from our results that they prefer genetically dissimilar males with whom they would produce offspring of low inbreeding coefficients. Females could increase the rate of mating with genetically dissimilar males by actively preferring dissimilar phenotypes (Gil et al. 2016) or by increasing the rate of spawning with increasing number of males joining a dominant male for multi-male spawning (Petersson and Jarvi 2001).

Salmonid fish are famous for their smelling ability (Keefer and Caudill 2014), and juvenile Arctic char learn to discriminate siblings by their MHC and further factors (Olsen et al. 2002). It is therefore possible that odors reveal alleles of the MHC via peptide ligands that are used as markers for the degree of kinship (Milinski et al. 2005). MHC-linked mate preferences could then be used to avoid genetically similar males, as has been found in a large number of other vertebrates, including various salmonids (Ruff et al. 2012). It remains to be tested whether MHC-linked mate preferences can lead to inbreeding avoidance in lake char and hence to the genetic benefits we observed (Landry et al. 2001).

In conclusion, we found that male yellowness does not predict offspring survival, growth, or stress resistance. This secondary sexual trait of lake char does therefore not seem to be an indicator of ‘good genes’. Females could, however, significantly increase offspring growth by avoiding genetically similar males with whom they would produce offspring that would suffer from high inbreeding coefficients. The potential genetic benefits of mate choice would be large if females aimed for such non-additive genetic effects (‘compatible genes’) instead of potential additive genetic effects (‘good genes’).

## Ethics

This work complied with the national, cantonal and university regulations where it was carried out. The handling and transport of adults and the transport of embryos were approved by the French authorities (INTRA.FR.2017.0109258).

## Data accessibility

All data have been deposited on the Dryad repository: https://datadryad.org/stash/share/t7cUjoEduKVxGR1AIlETlGXtqEFTrWXwBgEdHEKajcc https://doi.org/10.5061/dryad.gqnk98sp3 https://datadryad.org/stash/share/hsWb8yN3Km2jEUVa6_WvHTOpBJWC8050URzAnHQutLs

## Author contributions

LG, DN, CdG, LMC, EL, and CW designed the experiments. LG and EL organized the field work that was done by LG, DN, CdG, and LMC who also set up the experiments in the laboratory. LG was responsible for the 2nd experiment, DN for the 1st experiment, and CdG prepared the libraries, did the genomic analyses, and calculated inbreeding coefficients and kinships. LG, DN, LMC, LA, and AA monitored the embryos and determined growth. LG, DN, and CW did the statistics. CW wrote the manuscript that was revised by all authors.

## Acknowledgements

We thank G. Levray from the Pistolet fishery for catching the fish, L. Adhia Eya, A. Atherton, C. Berney, J. Bidaux, F. Dolivo, L. Espinat, S. Jacquet, S. Kreuter, E. Longange, J. Recher, and N. Sironi and for discussion and/or assistance in the field or in the lab, F. Schütz for discussion about statistics, and the Swiss SNF (31003A_182265) for financial support.

---

**Supplementary Figure S1.**
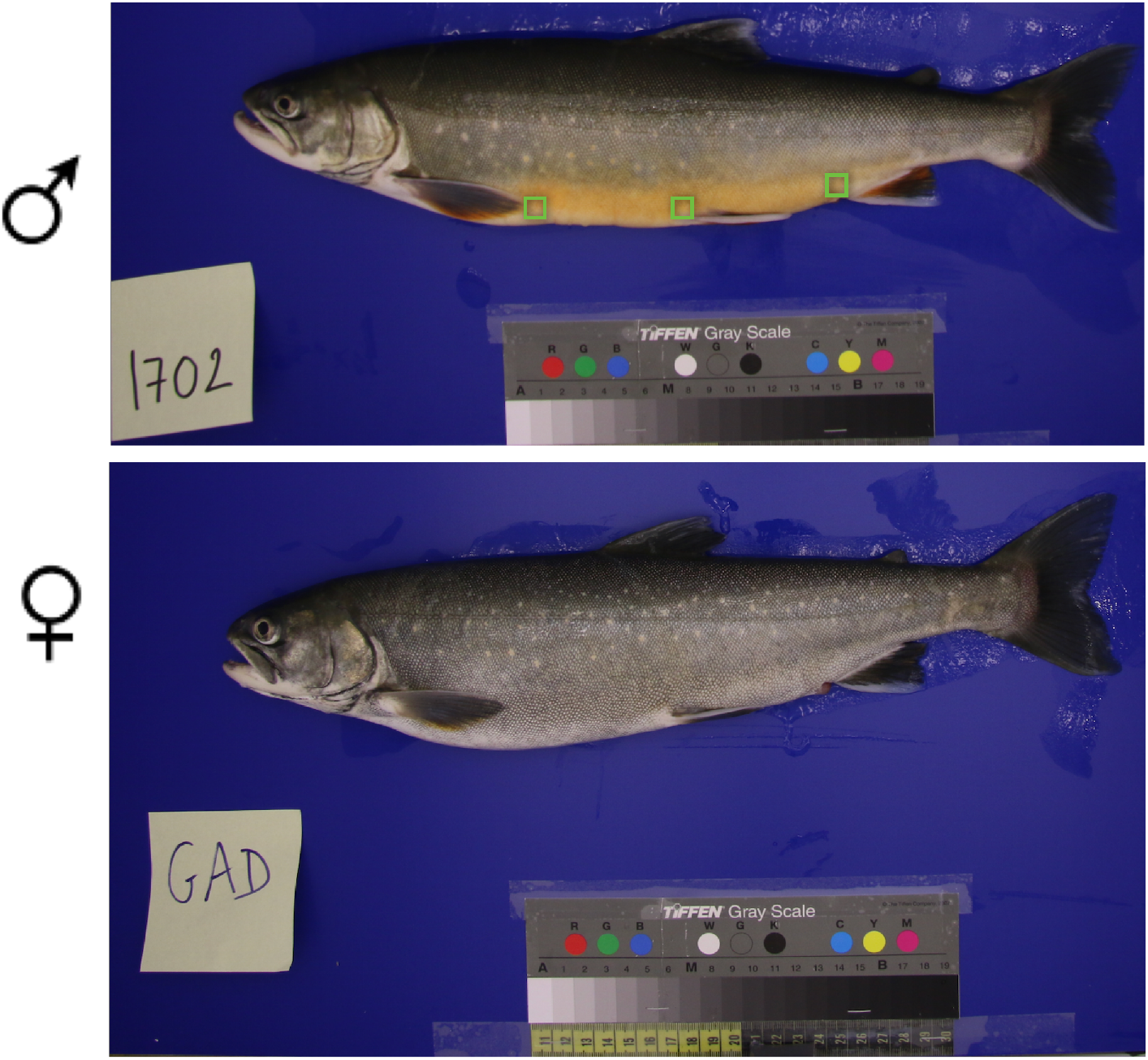
Size-standardized photos of a male of about median yellowness and a female lake char (*Salvelinus umbla*). The 3 green squares give the locations from which color measurements were taken. Photographs taken in a custom-made photobox under standardized light conditions (17 mm, f/5.0, 1/200 s, ISO 400, WB 4000 K, JPG 24 Mpx).

**Supplementary Figure S2.**
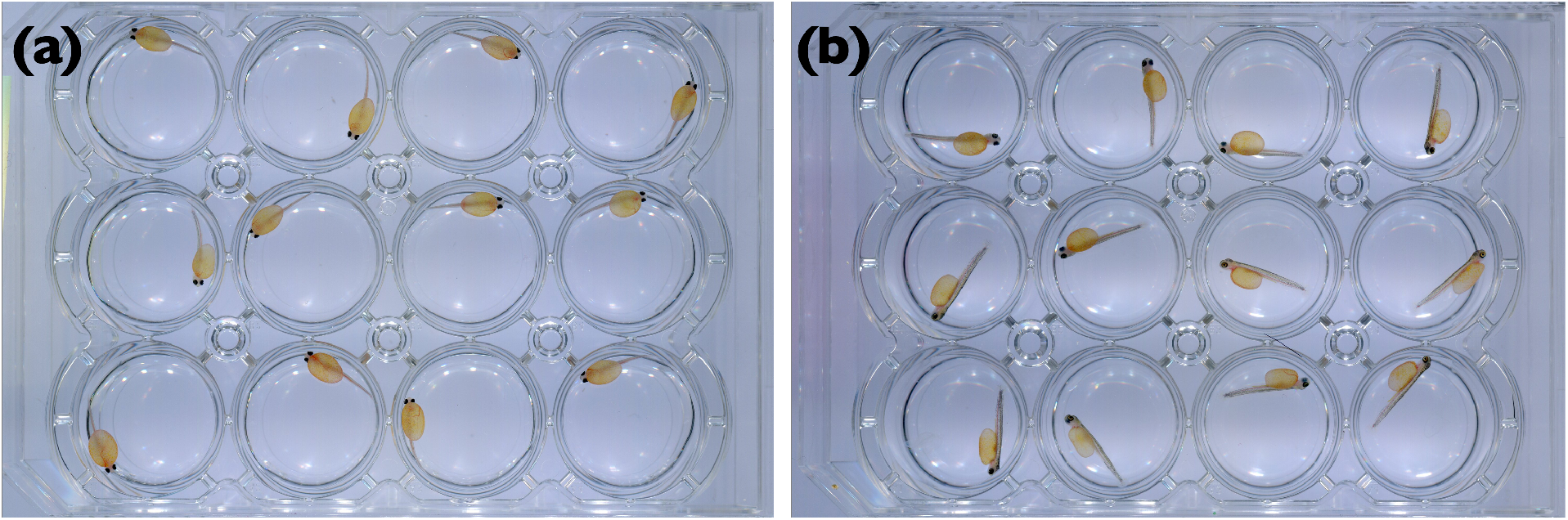
Yolk-sac larvae raised at 4.5°C in 12-well plates from hatching on. The photos were taken (a) at hatching and (b) 14 days later.

**Supplementary Figure S3.**
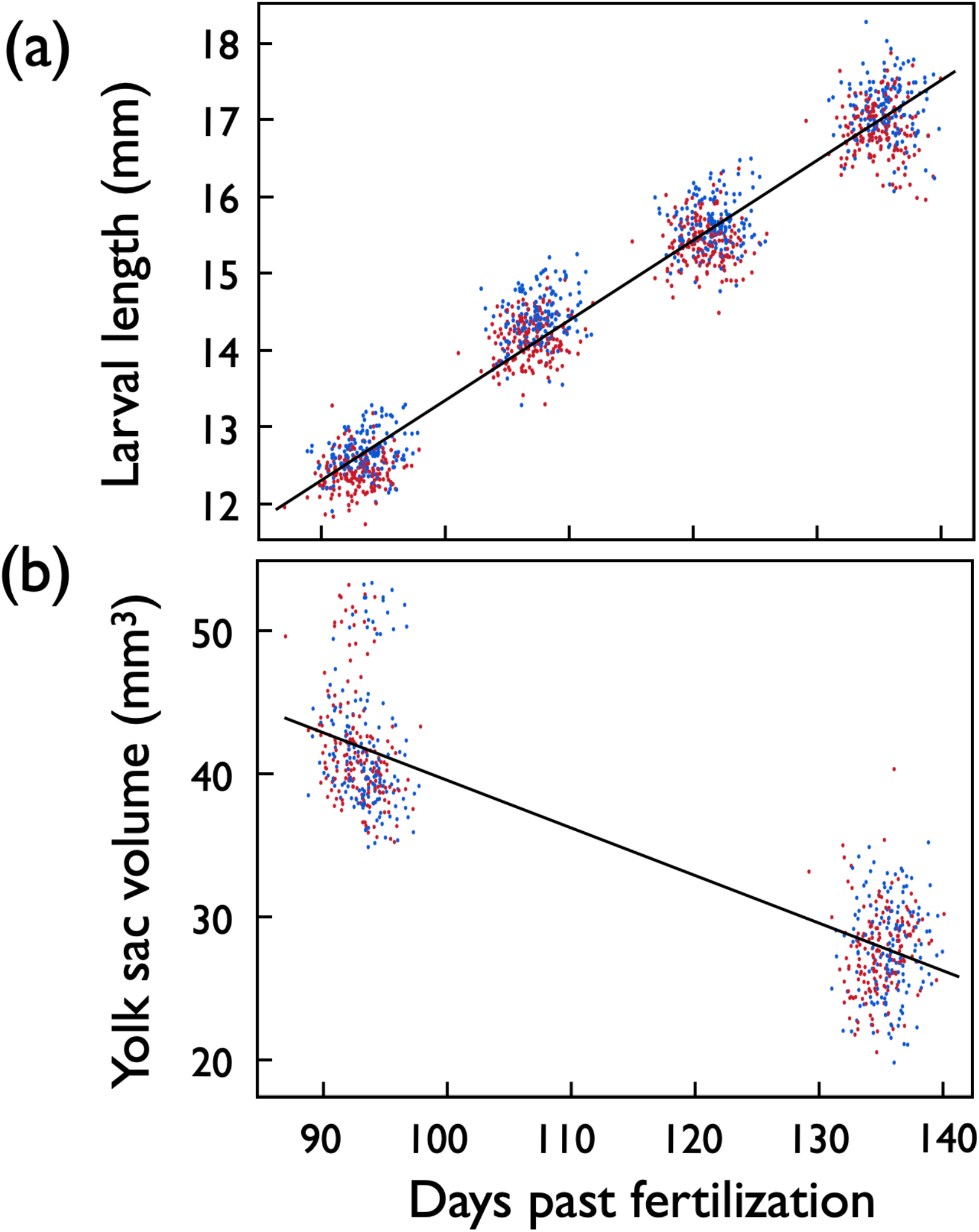
Days past fertilization (dpf) versus (a) average larval length and (b) average yolk sac volume per full-sib family and treatment (red: exposed to *Aeromonas salmonicida*, blue: sham exposed). Larval lengths were measured at the day of hatching and 14, 28, and 42 days later. Yolk sac volumes were measured at the day of hatching and 42 days later. The lines give the linear regressions.

**Supplementary Figure S4.**
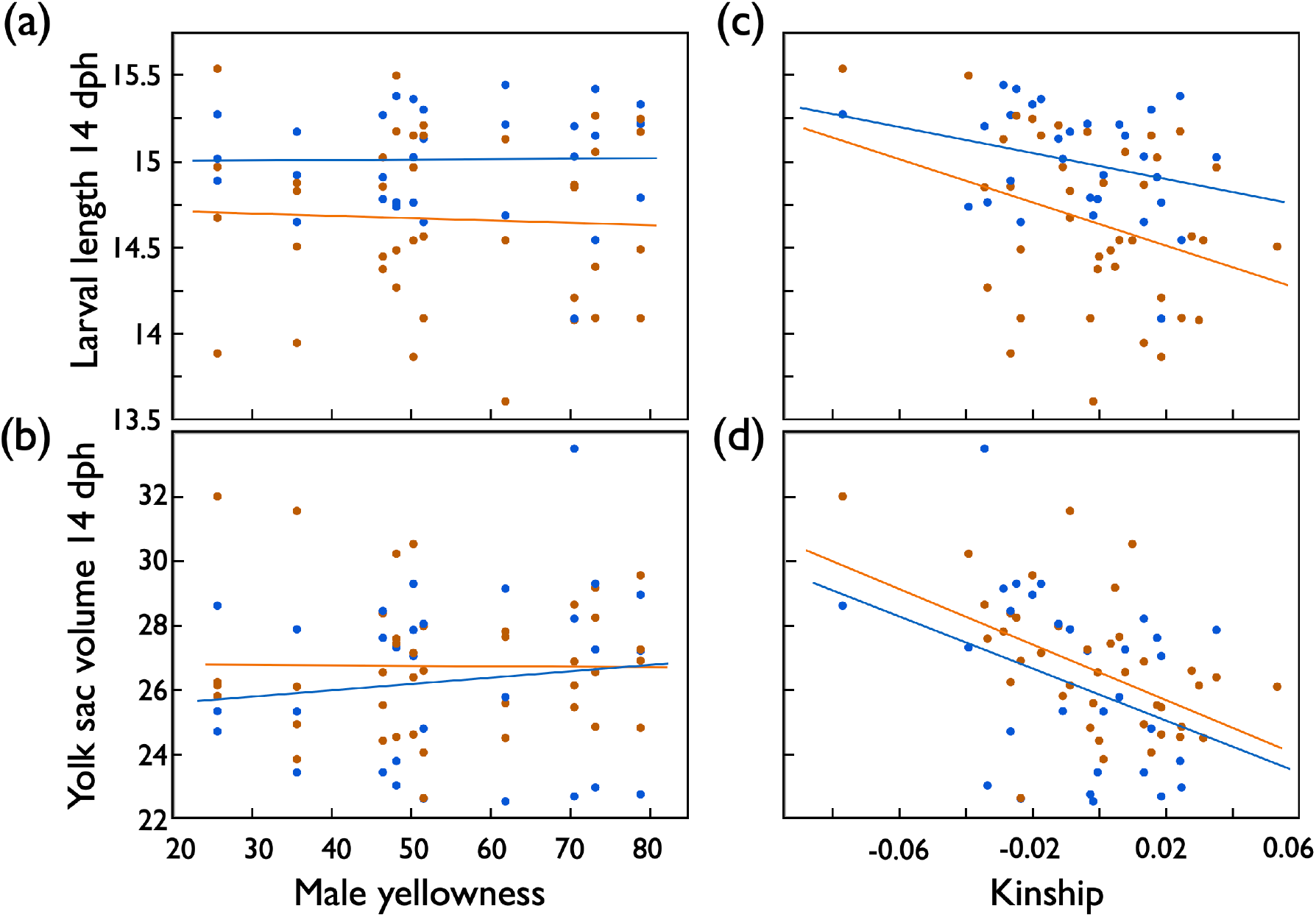
Larval growth in 1^st^ experiment: Mean larval length (mm) 14 days past hatching (dph), and mean yolk sac volume at that time (means per full-sib family) predicted by (a,b) male skin coloration (yellowness) or (c,d) by the genetic similarity between males and females (kinship) after exposure to remainders of ovarian fluids (orange symbols and regression lines), or sham exposed (blue). See Table S1 for statistics.

**Supplementary Figure S5.**
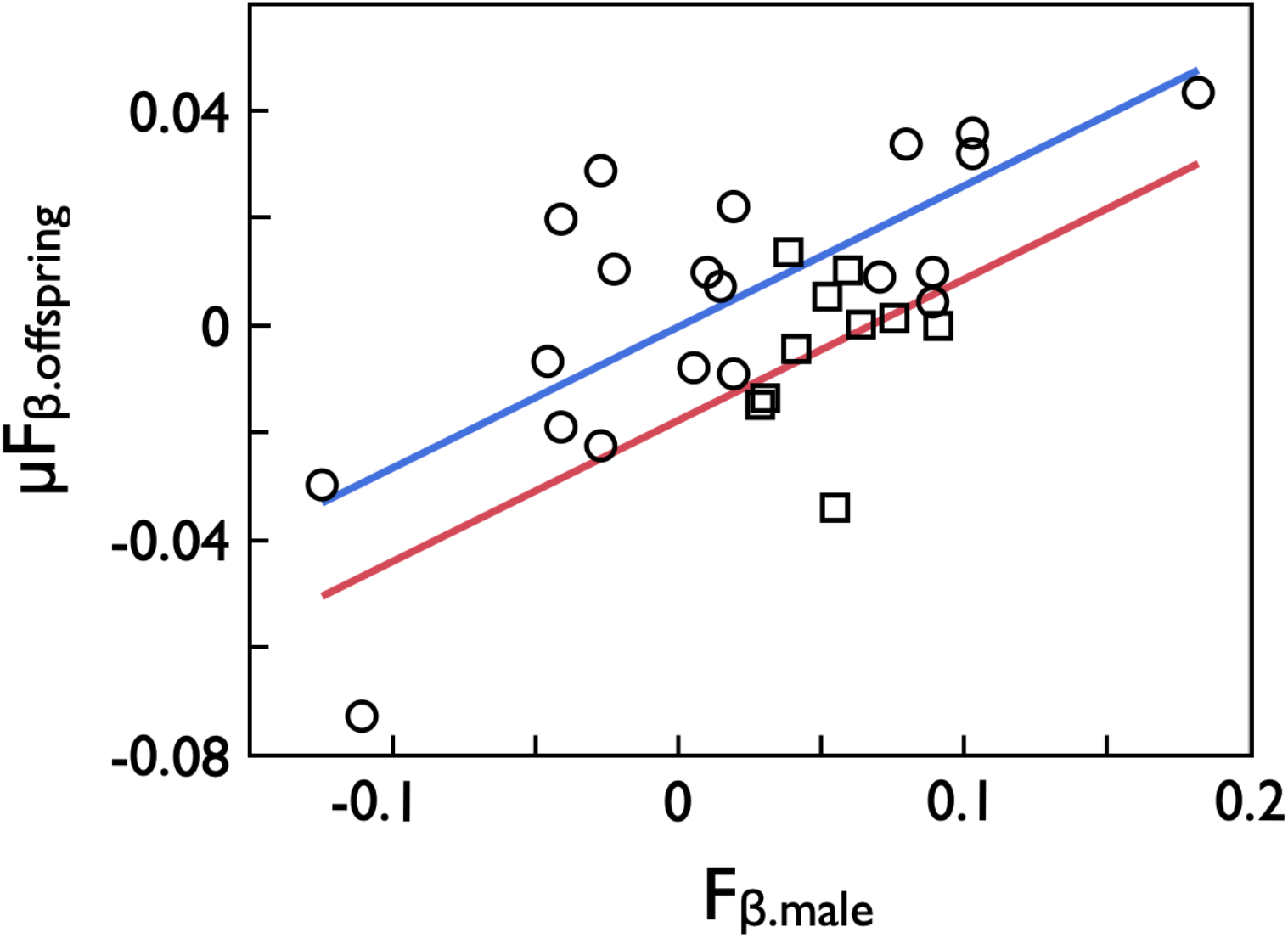
The mean expected inbreeding coefficients of each male’s offspring (µF_β.offspring_, corresponds to the mean kinship coefficients of each male to the females in the respective full-factorial breeding experiment) predicted by the males’ inbreeding coefficients (F_β.male_) for the 10 males of the 1^st^ experiment (squared symbols, red line) and the 21 males of the 2^nd^ experiment (round symbols, blue line). The lines give the fits from a multiple regression after removal of the non-significant interaction term. See text for statistics.

**Supplementary Table S1.** **Testing for effects on larval growth in 1^st^ experiment**: Linear mixed model on larval length 14 days post hatching (dph) when predicted by treatment and (a) male skin coloration (yellowness) or (b) the genetic similarity between males and females (kinship). Hatching date was included as further fixed factor. Family identity and the interaction between family ID and treatment were included as random factors. Significant p-values are highlighted in bold.

|  |  | Larval length 14 dph |  |  | Yolk sac volume 14 dph |  |  |
| --- | --- | --- | --- | --- | --- | --- | --- |
|  |  | Variance |  |  | Variance |  |  |
| Effects | d.f. | F | component | p | F | t | p |
| *(a) Male skin coloration* |  |  |  |  |  |  |  |
| Fixed effects: |  |  |  |  |  |  |  |
| Treatment | 1 | 17.1 |  | **<0.001** | 3.4 |  | 0.08 |
| Yellowness | 1 | 0.9 |  | 0.95 | 0.2 |  | 0.65 |
| Treatment x yellowness | 1 | 0.9 |  | 0.88 | 0.9 |  | 0.34 |
| Hatching date | 1 | 38.0 |  | **<0.001** | 79.5 |  | **<0.001** |
| Random effects: |  |  |  |  |  |  |  |
| Family ID<sup>1</sup> |  |  | 0.13±0.04 | **0.001** |  | 2.01±0.76 | **0.008** |
| Family x treatment<sup>1</sup> |  |  | 0.05±0.02 | **0.02** |  | 0.28±0.44 | 0.52 |
| Residual |  |  | 0.65±0.03 |  |  | 26.8±1.3 |  |
| *(b) Male & female kinship* |  |  |  |  |  |  |  |
| Fixed effects: |  |  |  |  |  |  |  |
| Treatment | 1 | 17.4 |  | **<0.001** | 4.5 |  | **0.04** |
| Kinship | 1 | 2.9 |  | 0.09 | 14.3 |  | **<0.001** |
| Treatment x kinship | 1 | 1.2 |  | 0.29 | 0.7 |  | 0.40 |
| Hatching date | 1 | 37.3 |  | **<0.001** | 86.9 |  | **<0.001** |
| Random effects: |  |  |  |  |  |  |  |
| Family ID<sup>1</sup> |  |  | 0.12±0.04 | **0.002** |  | 1.13±0.59 | **0.008** |
| Family x treatment<sup>1</sup> |  |  | 0.04±0.02 | **0.03** |  | 0.25±0.46 | 0.52 |
| Residual |  |  | 0.65±0.03 |  |  | 26.8±1.1 |  |

<sup>1</sup>REML unbounded variance components ± standard error, Wald p-values

